# A Bayesian psychophysics model of sense of agency

**DOI:** 10.1101/433888

**Authors:** Roberto Legaspi, Taro Toyoizumi

**Affiliations:** Laboratory for Neural Computation and Adaptation RIKEN Center for Brain Science RIKEN CBS-OMRON Collaboration Center 2-1 Hirosawa Wako City, Saitama 351-0198, Japan

## Abstract

Despite the increasing significance of sense of agency (SoA) research, the literature lacks a formal model: what computational principles underlie SoA, the registration that oneself initiated an action that caused something to happen? We theorize SoA in the framework of optimal Bayesian cue integration with mutually involved principles, namely, reliability of action and outcome sensory signals, their consistency with the causation of the outcome by the action, and the prior belief in causation. We used our Bayesian model to explain the intentional binding effect, hailed as reliable indicator of SoA. Our model explains temporal binding in both self-intended and unintentional actions suggesting that intentionality is not strictly necessary given high confidence in the action causing the outcome. Our Bayesian model also explains that if the sensory cues are reliable, SoA can emerge even for unintended actions. Our formal model therefore posits a *precision*-dependent *causal* agency.

---

The number of scientific contributions being added to the theoretical literature of sense of agency (SoA) has significantly increased at least in the past two decades^1,2^. The concept has garnered considerable attention in psychology, philosophy, neuroscience and psychopathology^3^. SoA is the registration^4^ that the self initiates an action in order to interact with and influence its external environment^5^. It has been posited that SoA is fundamental to the experience of volition^6-9^ and to self-consciousness because of its self-other distinction^10-12^, and the degradation of this experience characterizes certain psychiatric and neurological disorders^13-15^. Furthermore, SoA has recently been suggested to underpin neuroethics and law due to the role it plays in the social concept of responsibility for one’s own actions^5,9,16,17^.

Despite its increasing significance, the literature still lacks the computational principles that underlie SoA. We theorize SoA as the *confidence* in one’s perception of the action-outcome effect and that it is consistent (e.g., spatially or temporally) with the hypothesis that the action caused the outcome. We adapted the model of Sato, Toyoizumi and Aihara^18^ that was originally used to explain the *ventriloquism* effect as a Bayesian estimate of a common cause behind the consistency of the audiovisual stimuli. Formalizing SoA by this Bayesian psychophysics principle distinguishes our theory from existing works.

We compared the predictions of our model to the results of two pertinent intentional binding studies. Intentional binding, which is the perceived compression of the time interval between voluntary action and its outcome, has been reported as reliable implicit measure of SoA and has been used in a large number of studies providing valuable analyses on the temporal perception of action-outcome effects and the nature of SoA^19^. The seminal experiment of Haggard, Clark & Kalogeras^6^ investigated the perceived action-outcome timing effects in three conditions: voluntary wherein the subject intentionally presses a button, involuntary wherein muscle twitches of the subject’s hand are induced by a transcranial magnetic stimulation (TMS) applied to the motor cortex, and sham TMS wherein the TMS on the parietal cortex produces audible clicks but no movement (hereafter, voluntary, involuntary, and sham conditions, respectively). Haggard and colleagues computed the time interval between the perceived action timings (with the timings of either voluntary actions, muscle twitches, or audible TMS clicks as control experiment) and the perceived timings of subsequent tone stimuli. They showed that voluntary actions produced intentional binding, involuntary muscle twitches produced repulsion, i.e., prolonged opposite perception of the action-outcome intervals, and audible TMS clicks produced neither binding nor repulsion. Hence, they posit intentionality is necessary to achieve action-outcome binding.

The second pertains to the study of Wolpe, Haggard, Siebner & Rowe^20^ that investigated the contribution of cue integration to intentional binding by manipulating the reliability of the consequent tone relative to a background white noise. Such manipulation resulted in three levels of tone uncertainty conditions, namely, low, intermediate and high uncertainty. Their analyses showed that when tone reliability was reduced, the perceptual shift in tone timing towards the action was increased.

Our Bayesian model reproduces the above empirical results based on a computational principle. Further, it goes beyond timing estimations by exposing the underlying Bayesian mechanisms that possibly drove the temporal binding. Our Bayesian model explains the perceived compressed time interval between the action-outcome effect is more *consistent* with the prior belief of the *causal* role of one’s action in producing the immediate outcome, and thus increases the confidence in the Bayesian estimate assuming the causal case, modeled as SoA. Moreover, our model explains intentional binding as a specific class of the more general notion of causal binding. Our Bayesian model predicts that intentional binding generally happens on a per-trial basis, yielding a bimodal distribution of the perceived action-outcome interval. Lastly, the model also predicts that if the sensory input signals are perceived as reliable (precise), SoA may arise even for unintended actions, which serves as a testable theory for future SoA experiments. Our theory therefore provides a formal model that coherently accounts for the relationship between time perception, perception of causality, and reliability of action and outcome cues.

## Results

We considered the experimental setup of intentional binding where a subject presses a button (i.e., the action) and a tone (i.e., the outcome) sounds 250 ms after the button press. The true action and outcome timings are thus described by 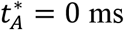 and 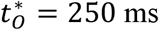, respectively, but they are unknown to the subject. The task for the subject is to accurately report her perceived timings of the button press and tone. We assume the arrival of relevant sensory input informing the timing of each of these physical events involves sensory delay *d* and jitter of variance σ^2^ due to sensory noise. Thus, the arrival time *τ*_*A*_ of sensory input that signals the action timing is assumed to be generated from a Gaussian distribution, 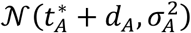, with mean 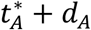 and variance 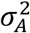. Similarly, the arrival time *τ*_*O*_ of sensory input that signals the outcome timing is generated from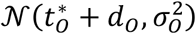.

The brain often resolves such ambiguity in sensory inputs by integrating multiple sensory cues akin to the Bayesian “ideal observer”^21^. Hence, we model a Bayesian observer who estimates action timing *t_A_* and outcome timing *t*_*O*_ based on the corresponding noisy sensory inputs arriving at time *τ*_*A*_ for the action and *τ*_*O*_ for the outcome. The conditional probability distributions of *τ*_*A*_ and *τ*_*O*_ that the Bayesian observer uses are modeled as Gaussian distributions

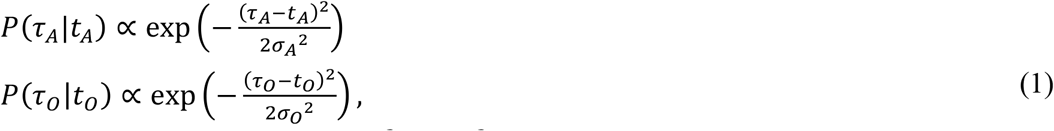

with mean *t_A_* and *t*_*O*_ and variance 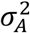 and 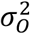 for action and outcome, respectively. Note that sensory delays *d_A_* and *d_O_* are not included in equation (1) for the reason we describe in the next paragraph.

Before studying the binding effect, let us consider simple baseline conditions. In one baseline condition, the action timing is reported by the subject without the presentation of an outcome tone. If no prior knowledge is available, the Bayesian observer reports the action timing that maximizes the conditional probability distribution in equation (1). Hence, the estimated action timing 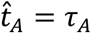 is solely determined by the noisy sensory input informing the action timing. In this case, the model predicts that the distribution of 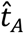 is 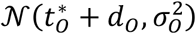. The mean and standard deviation of 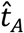 in the baseline condition were experimentally reported, e.g., Haggard’s results in the voluntary condition suggest *d_A_* = 6 ms and σ_*A*_ = 66 ms (refer to Table 1 in Methods for all condition-based *d_A_* and σ_*A*_ values). Importantly, we assume that the observer does not take into account sensory delay *d_A_* in equation (1). If the Bayesian observer included its effect, it could compensate for this delay and report unbiased timing, which was not the case in the experiment. Therefore, we assume that the observer was unable to take into account the sensory delay in equation (1). In the other baseline condition, the subject passively listens to a tone and reports its timing. This case goes parallel to the above case, and the model predicts that the estimated tone timing is 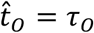, which is distributed according to 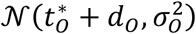. The comparison of this model prediction to Haggard’s experiment, for example, would be *d_O_* = 15 ms and σ_*O*_ = 72 ms (refer to Table 1).

**Table 1.**
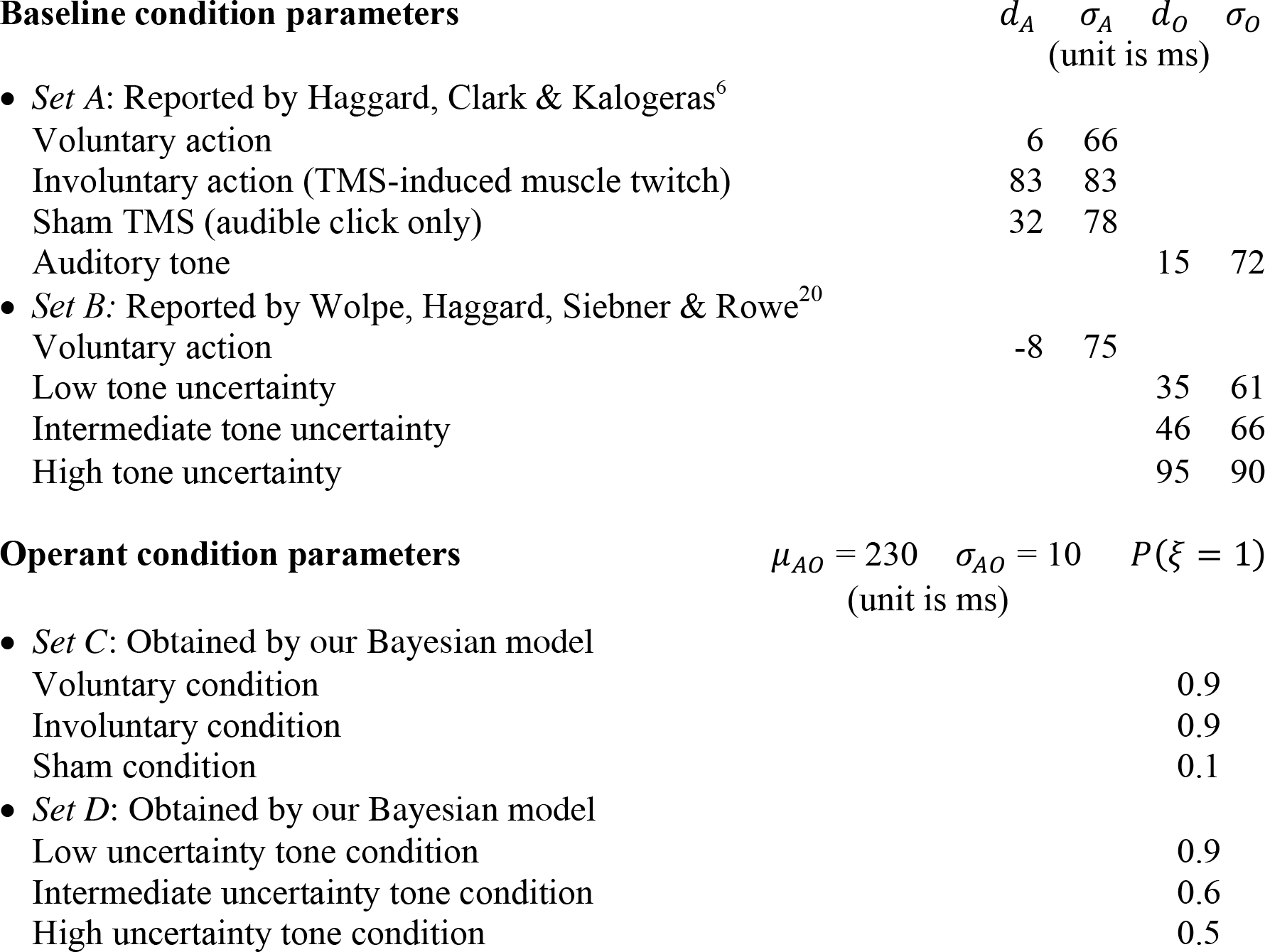
List of Bayesian model parameters and their values

Next, we study the effect of binding when the subject makes an action and then listens to the outcome tone, commonly referred to as the operant condition. In this case, the Bayesian observer makes an inference not only based on the conditional probability distribution in equation (1) but also based on the prior distribution of *t_A_* and *t*_*O*_. Adapting the Bayesian model of the v*entriloquism effect*^18^, we assume the prior distribution depends on the observer’s belief if the action caused the outcome, i.e., the *causal* case: ξ = 1, or the action and the outcome are unrelated, i.e., the *acausal* case: ξ = 0:

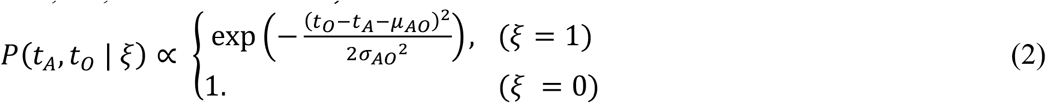

The action causes the outcome in the causal case (*ξ* = 1) so that the outcome timing involves a typical delay *μ*_*AO*_ with respect to the action timing and a Gaussian-distributed jitter of standard deviation σ_*AO*_. The outcome is caused by something other than the action in the acausal case (*ξ* = 0) so that *t_A_* and *t*_*O*_ are independent. Lastly, we define *P*(*ξ*) as the *prior* for each belief: *P*(*ξ* = 1) for the causal case and *P*(*ξ* = 0) = 1 − *P*(*ξ* = 1) for the acausal case. We hypothesize the estimation of *ξ* to be essential for the perception of causality and SoA (explained below).

The prior probability of equation (2) cannot be normalized unless a finite range of (*t_A_*, *t*_*O*_) is defined. Therefore, we only consider it in the range 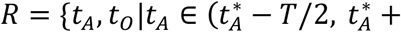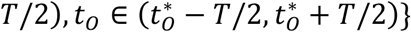 and assume that it is zero outside *R*, where again 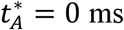 and 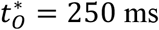 are the true action and outcome timings, unknown to the observer, and *T* = 250 ms is a large enough but finite constant that specify the interval lengths in consideration. Hence, the prior probability distribution *P*(*t_A_*, *t*_*O*_ | *ξ*) in equation (2) must be normalized within *R*. Our results are robust to a shift in the center of *R*.

Given a pair of sensory inputs at *τ*_*A*_ and *τ*_*O*_, the Bayesian observer estimates the most probable timing for the action and the outcome and whether these observations are consistent with the causal case. According to the Bayesian estimation theorem, the maximum-a-posteriori (MAP) estimate 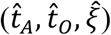 of the corresponding pair of physical sensory timing (*t_A_*, *t*_*O*_) and the causal variable *ξ* is given by

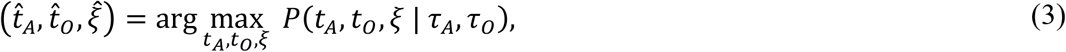

where *P*(*t_A_*, *t*_*O*_, *ξ* | *τ*_A_, *τ*_*O*_) is the *posterior* probability distribution of (*τ*_*A*_, *τ*_*O*_, *ξ*) given the sensory inputs (*τ*_*A*_, *τ*_*O*_). Hence, whether the Bayesian observer estimates the action-outcome effect to be causal or not depends on the posterior-ratio comparing the causal case (*ξ* = 1) and the acausal case (*ξ* = 0), namely,

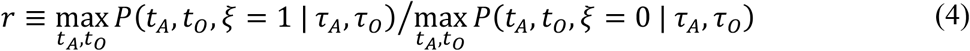

Causality is detected if the confidence in the causal estimate is greater than that in the acausal case, that is, *r* > 1. The MAP estimate of equation (3) is then given by (see Methods for the derivation)

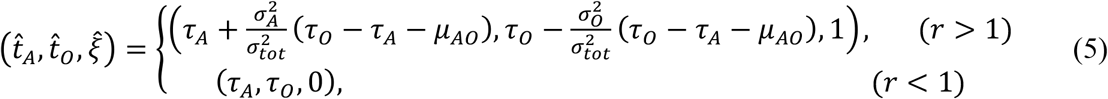

with 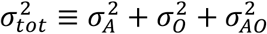. This indicates, on one hand, that perceptual shift does not happen if the causality is not detected 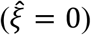 − the time estimates for action and outcome simply reflect the corresponding sensory signals in this case. On the other hand, perceptual shift happens if the causality is detected 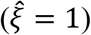 − the action and outcome timing attract each other in the form of binding if *τ_O_* − *τ_A_* > *μ_AO_* and repel each other in the form of repulsion if *τ_O_* − *τ_A_* < *μ_AO_*. The magnitude of perceptual shift for the action and outcome timing is given by 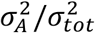 and 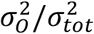, respectively, implying that perceptual shift is greater for a more unreliable stimulus. This model predicts that the occurrence of binding, repulsion, or no perceptual shift is trial-dependent, influenced by the noisy sensory signal *τ_O_* − *τ_A_* informing the action-outcome interval. We denote the probability of detecting causality (i.e., 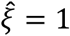) by *P_c_* (see Method for its analytical expression). *P_C_* increases with larger *P*(*ξ* = 1) and smaller σ_*AO*_ if σ_*AO*_ << σ_*A*_,σ_*O*_.

Lastly, separately from the judgement of causality described above, we also directly quantify the confidence in the causal MAP estimate

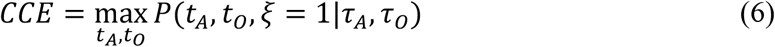

and we postulate this quantity to be a possible indication of the pre-reflective feeling of agency (see Discussion). The analytical expression of *CCE* in the appendix yields the following requirements to have high *CCE*: (A) The timing of sensory signals must be consistent with the causation of the outcome by the action, namely, *τ*_*O*_ − *τ*_*A*_ ≈ *μ_AO_* (B) The causal prior probability *P*(*ξ* = 1) must be high; (C) The sensory inputs must be precise, i.e., the amplitudes σ_*A*_ and σ_*O*_ of sensory jitter must be small enough. We therefore posit SoA as encapsulation and manifestation of several pertinent aspects, which include temporal consistency in the action-outcome effect, the prior belief of an action causing the outcome, and the reliability of the perceived sensory signals.

Here, we briefly describe how we obtained the parameter values used in our simulation (but see Methods for more details about the simulation and model fitting). Fitting of *d_A_*,*d_O_*, σ_*A*_ and σ_*O*_ is straightforward, they are suggested by the means and standard deviations of the reported subjects’ baseline estimation errors (Table 1-Sets A and B). After fixing these parameters, the model is left with three free parameters, *μ*_*AO*_, *σ*_*AO*_, and *P*(*ξ* = 1). As described in equation (5), *μ*_*AO*_ has an important role in determining if binding or repulsion happens in each experimental condition. A fixed value of *μ*_*AO*_ = 230 ms successfully accounts for this qualitative behavior in all the 6 experimental conditions (3 from Haggard et al. and 3 from Wolpe et al.) that we study. The analytical expressions in Methods suggest that *σ*_*AO*_ and *P*(*ξ* = 1) have a largely overlapping role in detecting causality. Causality is more likely detected if *σ*_*AO*_ is small or *P*(*ξ* = 1) is large, although the exact mechanisms are slightly different. At least one of these two parameters needs to be adjusted according to the conditions to account for the experimental observations. For simplicity, we fix σ_*AO*_ = 10 ms to be a small enough constant to permit noticeable perceptual shift and adjust *P*(*ξ* = 1) (see Table 1 for the parameter values in 6 experimental conditions) to account for two observations in each condition, namely, the perceptual shifts in the action timing and the outcome timing.

Our results show that our simple Bayesian model qualitatively reproduces the perceptual shifts that were reported in Haggard et al.’s study (Fig. 1). Consistent to their findings, our Bayesian observer inferred the action and outcome perceptual shifts to bind in the voluntary condition resulting to compressed temporal intervals between the action and outcome perceptual shifts. However, repulsion, i.e., reversed and prolonged perceptual shifts, was observed for the involuntary condition. The model also reproduced no appreciable perceptual shifts in the sham condition.

**Figure 1.**
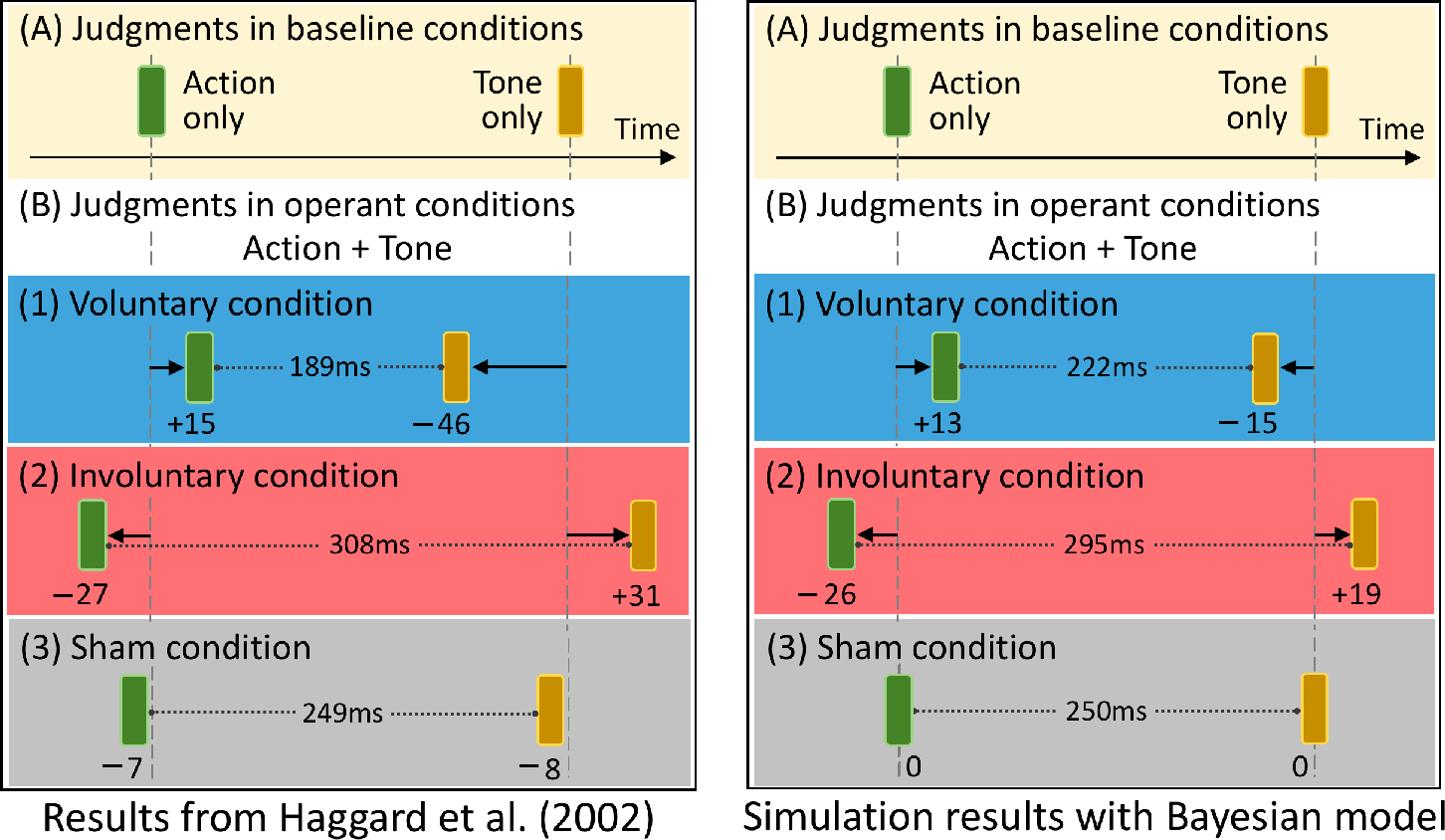
The qualitative replication of the empirical results reported by Haggard, Clark & Kalogeras^6^ (left panel) by our Bayesian model (right panel). The subjects’ mean judgment error for the single-event baseline condition is subtracted from the mean judgment error for the corresponding operant event, which results to the magnitude and direction of the perceptual shifts. A *positive* perceptual shift informs delayed awareness, and a *negative* shift informs anticipated awareness. The action and outcome timings are perceived to shift towards each other in the voluntary condition. In contrast, they are perceived to repulse in the involuntary condition. There is no discernible perceptual shift in the sham condition.

Our Bayesian model predicts binding and repulsion to increase with stronger causal prior (Fig. 2). From equation (5), the perceptual binding in the action-outcome interval is given by 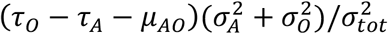 in the causal case 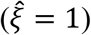 and none otherwise 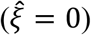. Because the sensory signals are distributed according to 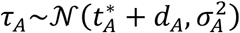 and 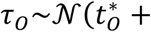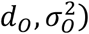, the average of *τ*_*O*_ − *τ*_*A*_ − *μ*_*AO*_ factor is 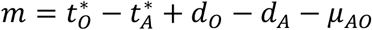. Hence, the sign of *m* determines if binding or repulsion is predicted on average. With the current set of parameters, *m* is positive in the voluntary condition, yielding binding, and negative in the involuntary condition, yielding repulsion (schematically drawn in Fig. 3a). Perceptual shift is almost zero regardless of the causal prior *P*(*ξ* = 1) in the sham condition because *m* ≈ 0. We chose *P*(*ξ* = 1) = 0.1 for this under-constrained sham condition, assuming that causality would not be frequently detected.

**Figure 2.**
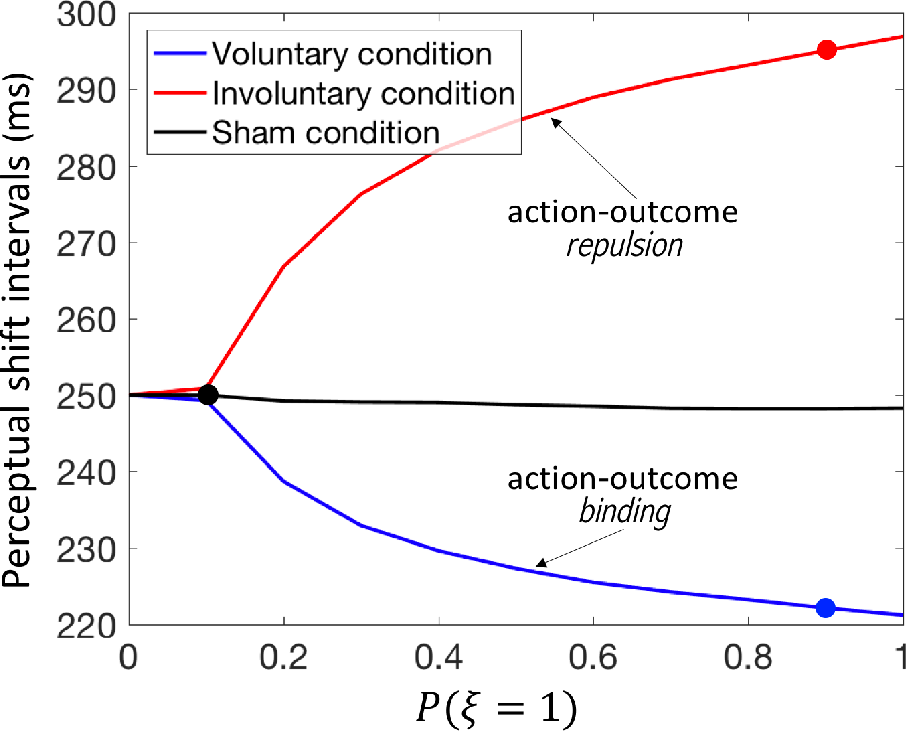
The Bayesian model predictions of the influence of causal prior strength on action-outcome perceptual shifts. The best estimates of the Bayesian model (in Fig. 1) were obtained from different causal priors, specifically, *P*(*ξ* = 1) is 0.9, 0.9 and 0.1 (marked by the colored dots) for the voluntary, involuntary, and sham conditions, respectively. The intervals between the action and outcome perceptual shifts shrink in the voluntary, but widen in the involuntary, condition with a strong causal prior. Minimal changes in perceptual shifts are predicted for the sham condition even with a strong causal prior.

Our Bayesian model provide interesting insights on what possibly drives the perceived action-outcome temporal binding and repulsion effects. We empirically observed sensory delay *d* to increase with larger standard deviation σ of the Gaussian-distributed jitter (observed in both Haggard and Wolpe; see Table 1). This may imply that, as action or outcome ambiguity is increased due to noise (greater σ) for increased sensory uncertainty, more time would be needed (greater *d*) for a sensory input to reach the subject’s perceptual threshold for temporal awareness in the baseline condition. Thus, because of the *m*’s dependency on *d*_*O*_ − *d*_*A*_, binding more likely happens when the outcome is unreliable (i.e., with large *d*_*O*_) and repulsion more likely happens when the action is unreliable (i.e., with large *d*_*A*_).

To further illustrate the model prediction from our simulations, we plotted separately the action and outcome perceptual shifts for the voluntary, involuntary and sham conditions as functions of the temporal disparity *τ*_*O*_ − *τ*_*A*_ (c.f. equation (5)). Indeed, our data shows that for instances in which *τ*_*O*_ − *τ*_*A*_ > *μ_AO_*, action awareness is delayed (positive action shift, Fig. 3b) and outcome tone is anticipated (negative outcome shift, Fig. 3c), thereby demonstrating binding. The opposite happens when *τ*_*O*_ − *τ*_*A*_ < *μ_AO_* thereby demonstrating repulsion in both action and outcome awareness (Figs. 3b and 3c, respectively). We then plotted how the model’s MAP estimates on the action-outcome interval are affected by the sensory time difference *τ*_*O*_ − *τ*_*A*_ in the baseline (here 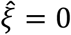 is forced; Fig. 3d) and operant (Fig. 3e) conditions. We observe from the baseline condition that the MAP estimates follow sensory inputs, 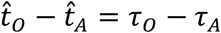, whereas the perception of action and outcome timings shifted towards the prior mean, 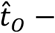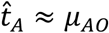, in the voluntary and involuntary conditions but not so much in the sham condition with weak causal prior. Therefore, our model is agnostic as to whether the action is self-intended or unintended. Binding towards 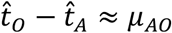 will happen, be it in the opposite direction, as long as the action is believed to have caused the outcome. This suggests that causality is the phenomenon that underlies temporal binding, and likely SoA, with self-intended causality being a specific case. The temporal window of *τ*_*O*_ − *τ*_*A*_ for detecting causality is wider in the voluntary and involuntary conditions than the sham condition.

**Figure 3.**
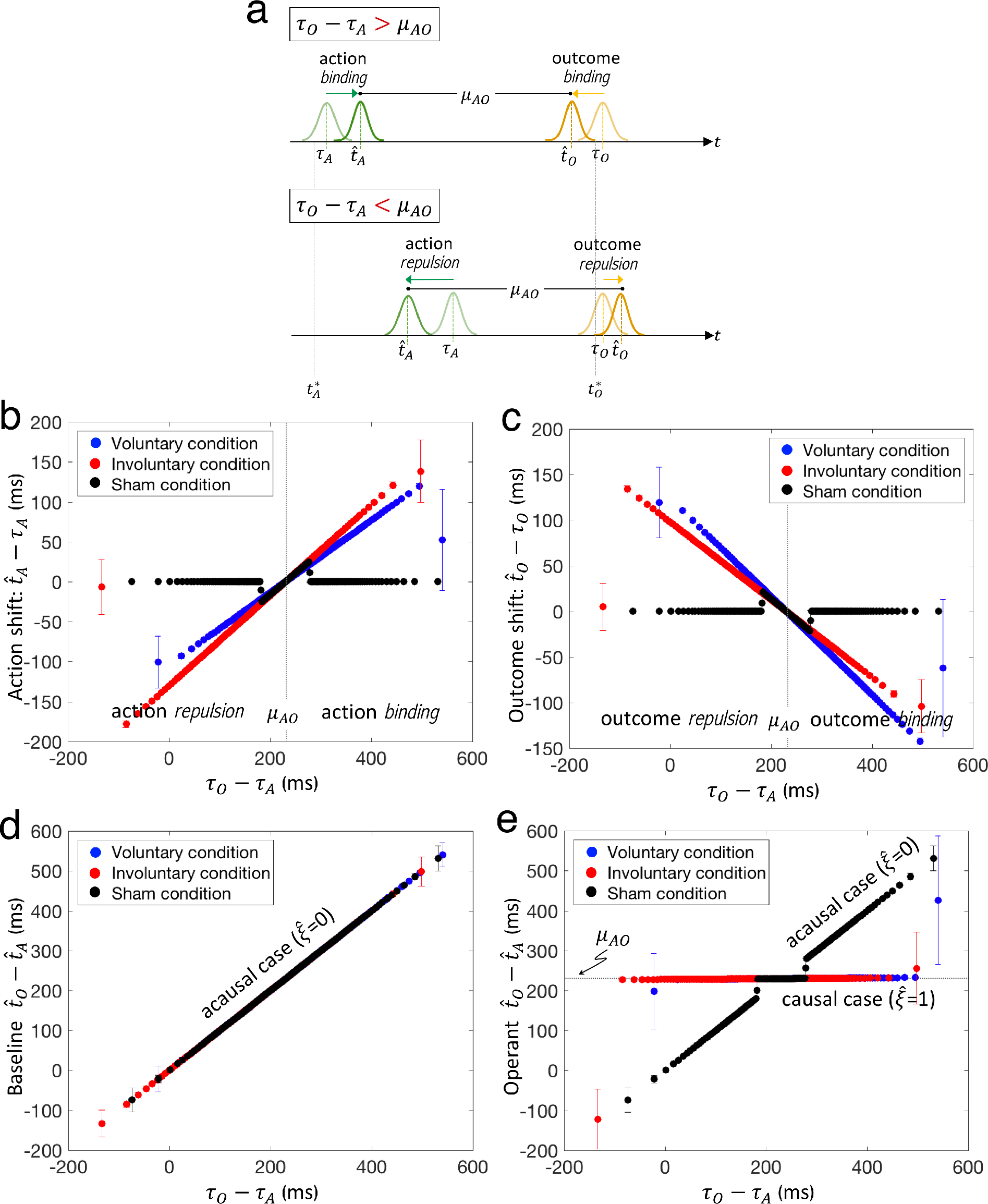
The Bayesian model predictions of trial-to-trial action-outcome temporal binding and repulsion effects, in **(b)** and **(c)**, and action-outcome timing interval, 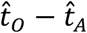, in the baseline and operant conditions, in **(d)** and **(e)**, as functions of the perceived temporal disparity, *τ*_*O*_ − *τ*_*A*_. The causal prior *P*(*ξ* = 1) is 0.9, 0.9 and 0.1 for the voluntary, involuntary, and sham conditions, respectively (as in Fig. 2). The per trial results are grouped accordingly into bins of width 200 (randomly chosen), and the mean and standard deviation for each bin are plotted. This format is followed each time a quantity of interest is plotted as a function of *τ*_*O*_ − *τ*_*A*_. **a|** Our Bayesian model predicts that (shown schematically) if *τ*_*O*_ − *τ*_*A*_ > *μ*_*AO*_ action and outcome binding will happen. Otherwise, i.e., *τ*_*O*_ − *τ*_*A*_ < *μ*_*AO*_, action-outcome repulsion will occur. In both cases, the perceived timings in the baseline move (compress or stretch) towards the temporal consistency 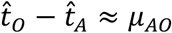 in the operant condition. **b, c|** When *τ*_*O*_ − *τ*_*A*_ > *μ*_*AO*_, there is positive perceptual shift in action awareness 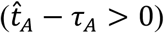 and negative perceptual shift in outcome awareness 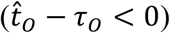. The opposite happens when *τ*_*O*_ − *τ*_*A*_ < *μ*_*AO*_. *Both* binding and repulsion occur in *both* voluntary and involuntary conditions, but very little effect in the sham condition. **d|** The Bayesian estimates follow the sensory inputs in the baseline condition, i.e., 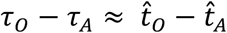, where all trials are acausal 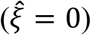 by definition. **e|** The Bayesian estimate shifts toward the prior assumption, 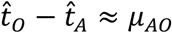, when the sensory inputs are highly consistent with the prior, *τ*_*O*_ − *τ*_*A*_ ≈ *μ*_*AO*_, and therefore when causality is detected 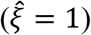. Otherwise, the estimate of action and outcome timings follow the sensory inputs.

We then examined how the prior belief in causation affects our proposed measure for SoA in Haggard’s experimental setup. Our model predicts *CCE* to strengthen together with the causal prior but its strength differs depending on the conditions even at the same strength of the prior (Fig. 4a). Interestingly, *d_A_* and σ_*A*_ are the only parameters of our Bayesian model that differentiate the three conditions in this figure. As we described above, these two parameters are empirically correlated such that the delay *d_A_* increases with larger σ_*A*_. Hence, the difference in *CCE* in the three conditions can be attributed to the inequalities in the standard deviations of the subjects’ action timing estimation errors in the three conditions: 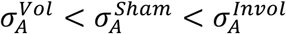 as per Haggard et al.’s data. Haggard et al. speculated that the unexpected and surprising quality of the TMS-induced movement could account for the repulsion effect in the involuntary condition. We suggest that this surprise might have introduced uncertainty in the perception of action input signals. Hence, while subjects were certain of the nature of their voluntary actions, they could be less certain of the proprioception signals induced by TMS, which could explain the inequalities in σ_*A*_. As a result, the model gives *CCE^Vol^* > *CCE^Sham^* > *CCE^Invol^* according to the requirement (C), i.e., reliable sensory inputs, for having high *CCE* when compared at the same strength of the causal prior.

**Figure 4.**
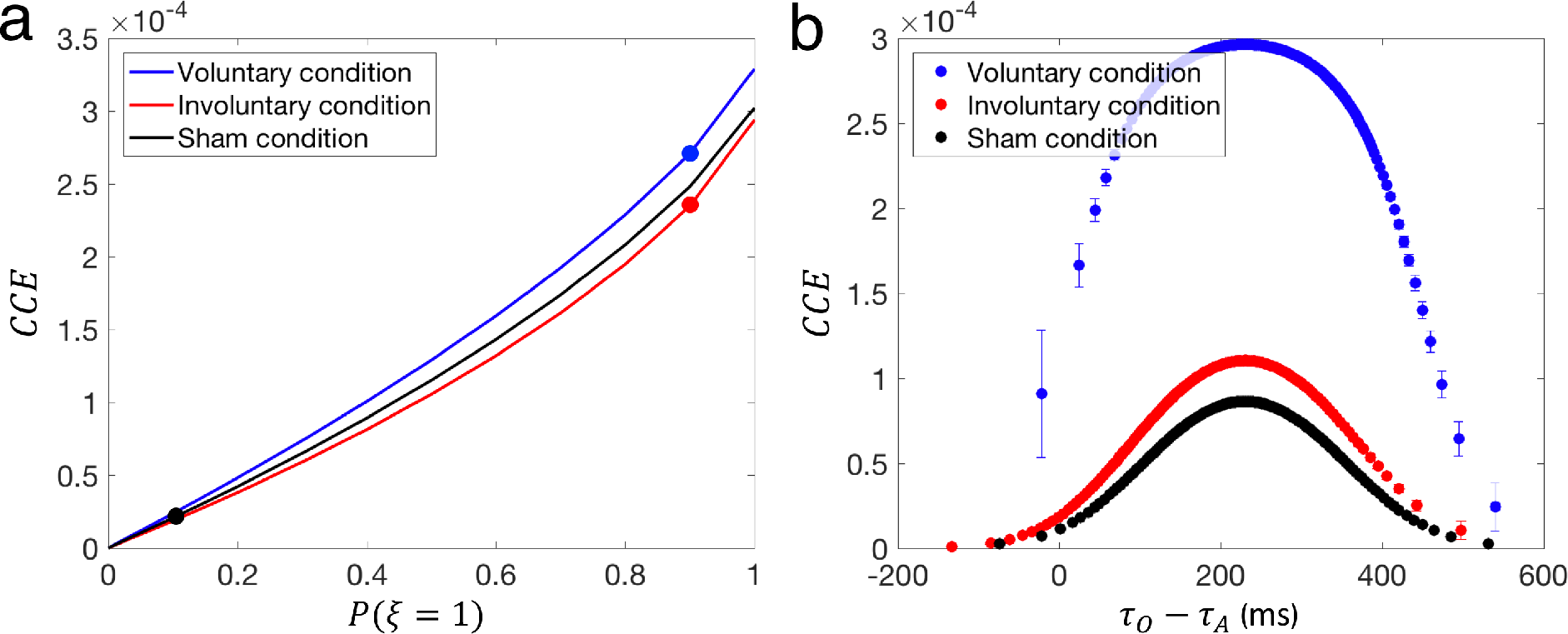
The Bayesian model predictions of *CCE*, i.e., the *confidence in causal estimate*, which is our proposed measure for SoA. **a**| Our Bayesian model predicts *CCE* to increase with a stronger causal prior. Furthermore, *CCE* differs for each condition even with equal prior strengths. This can be attributed to the difference in the amplitude of the jitter in the self-generated versus TMS-induced (muscle twitches and audible clicks) movement. **b**| When plotted as functions of the trial-to-trial temporal disparity *τ*_*O*_ − *τ*_*A*_, with the specific causal priors obtained for each condition, marked in **(a)**, *CCE* has a higher peak in the voluntary condition, but much lower values in the sham condition. Furthermore, *CCE* diminishes as the temporal disparity in sensory inputs moves further away from the prior mean, |*τ*_*O*_ − *τ*_*A*_ − *μ*_*AO*_. This falling of the *CCE* is faster when the causal prior is weaker and the uncertainty in the action input signal is higher.

The relation between *CCE* and SoA becomes clear when we analyze them with the fitted values of the causal prior (*P*(*ξ* = 1) = 0.9 for the voluntary and involuntary conditions and *P*(*ξ* = 1) = 0.1 for the sham condition as indicated in Table 1). Figure 4b plots *CCE* on a per trial basis as functions of the temporal disparity *τ*_*O*_ − *τ*_*A*_ (c.f. the analytical expression for *CCE* in Methods). *CCE* in the voluntary condition has a higher peak than the involuntary condition as we described above (due to small σ_*A*_ in the voluntary condition for the requirement (C)). In both voluntary and involuntary conditions, *CCE* diminishes as *τ*_*O*_ − *τ*_*A*_ moves farther from *μ*_*AO*_ because of the requirement (A) of small |*τ*_*O*_ − *τ*_*A*_ − *μ*_*AO*_| for having high *CCE*. Finally, *CCE* for the sham condition takes much lower values than the voluntary or involuntary conditions because of the requirement (B) of large *P*(*ξ* = 1) for having high *CCE*. Hence, our Bayesian model coherently explains not just SoA that arises from the causation of the outcome by the action, but also the one that is influenced by the reliability of the different agency cues − a *precision*-dependent *causal agency*.

In a similar fashion, we then examined the underlying psychophysical mechanisms that could account for the temporal binding observed by Wolpe and colleagues, in which three uncertainty levels (high, intermediate, and low uncertainty) of the outcome stimulus were tested. We use the Bayesian model that was used to reproduce the Haggard’s experiments with the same values of *μ*_*AO*_ and *σ*_*AO*_ but adjusted the strength of the causal prior *P*(*ξ* = 1) to fit the reported action timing and outcome timing in each condition. We used *P*(*ξ* = 1) = 0.9, 0.6 and 0.5 for low, intermediate and high tone uncertainty conditions, respectively (see Table 1 and Methods). This means that the prior belief in causation decreases with the tone uncertainty, which is plausible.

Our model reproduces the experiments of Wolpe et al. (Fig. 5a), qualitatively explaining the temporal binding they observed in terms of a single, coherent cue integration formulation. The Bayesian estimate of the action-outcome intervals shift towards 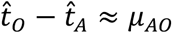, as per the causal temporal prior in equation (2) when causality is detected. On the one hand, the magnitude of the shift is greater when the outcome uncertainty is high (c.f. equation (5)). But, on the other hand, causality is less frequently detected when the outcome uncertainty is high with the reduced causal prior. These two opposing effects are summarized in Fig. 5b. The model can qualitatively reproduce the experiments if the former effect is more dominant. Quantitatively, however, the latter effect is necessary to mitigate the former effect.

**Figure 5.**
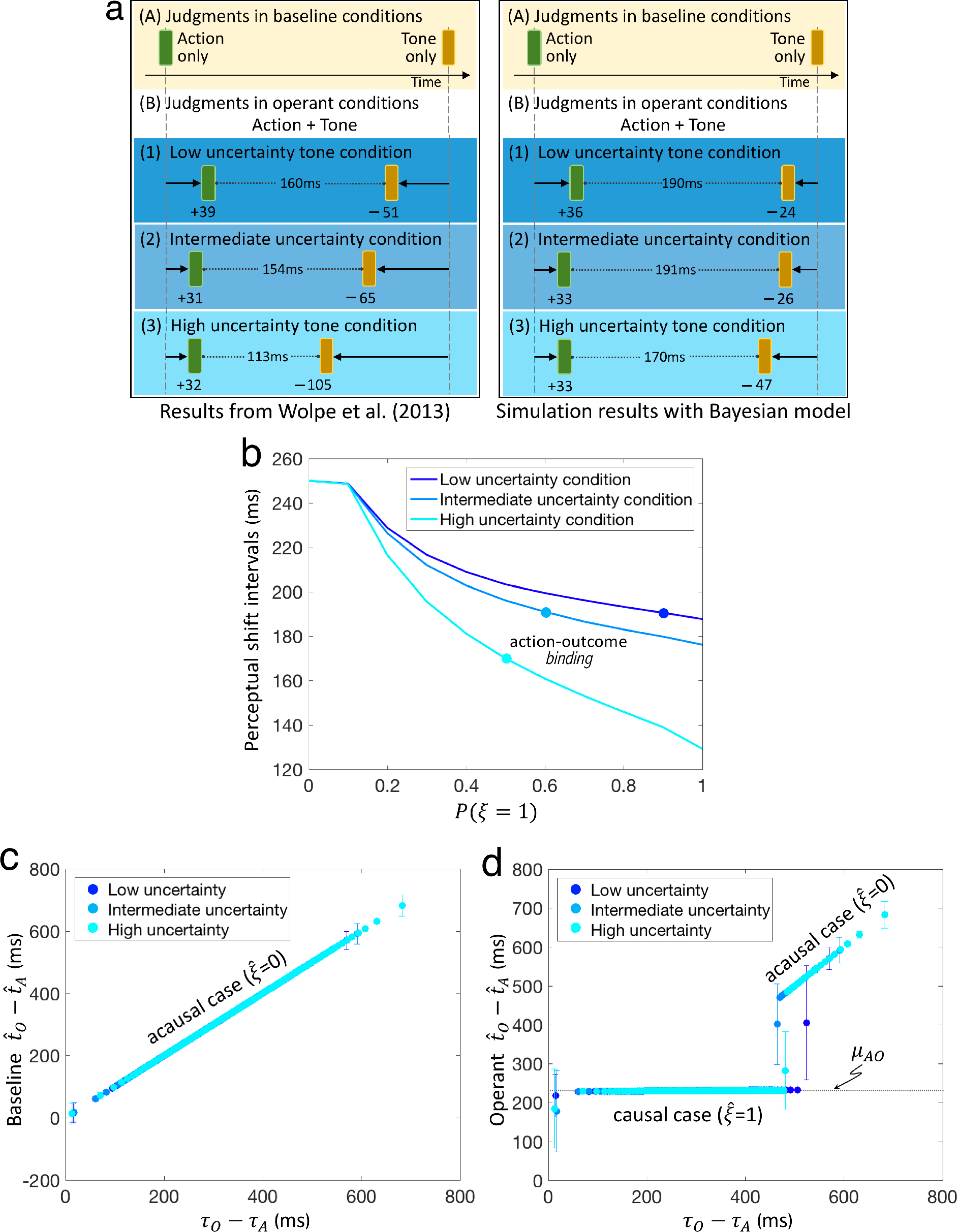
The Bayesian model qualitative replications of, as well as predictions related to, the results reported by Wolpe, Haggard, Siebner & Rowe^20^. **a|** Qualitative replication of the experimental results (left panel) by our Bayesian model (right panel). **b**| The action-outcome binding increases under heightened uncertainty. However, causality is less detected when the causal prior is lower, which decreases the action-outcome binding effect. The best estimates of the Bayesian model in **(a)** were obtained from different causal prior strengths, specifically, *P*(*ξ* = 1) is 0.9, 0.6 and 0.5 (marked by the colored dots) for the low, intermediate and high tone uncertainty conditions, respectively. **c, d**| The causal prior strengths that correspond to each condition were used for the Bayesian estimate of the action-outcome timing interval 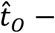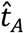 in the baseline and operant conditions. The Bayesian estimate follows the sensory inputs in the baseline condition where all trials are acausal, but shifts toward the prior assumption, *τ*_*O*_ − *τ*_*A*_ ≈ *μ*_*AO*_, when causality is detected. The temporal window of *τ*_*O*_ − *τ*_*A*_ for detecting causality is wider when the outcome uncertainty is lower, which means more instances demonstrate binding.

Next, we plot how the Bayesian estimate of the action-outcome interval, 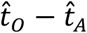, depends on the sensory inputs, *τ*_*O*_ − *τ*_*A*_. The perceived intervals faithfully follow the sensory inputs in the baseline condition (Fig. 5c), where all trials are acausal 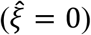 by definition. In the operant condition (Fig. 5d), the Bayesian estimate shifts toward the prior assumption 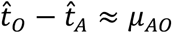 when the sensory inputs are highly consistent with the prior *τ*_*O*_ − *τ*_*A*_ ≈ *μ*_*AO*_ and, thus, when the causality is detected 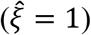. Otherwise, the estimate of action-outcome intervals follows sensory inputs. The temporal window of *τ*_*O*_ − *τ*_*A*_ for detecting causality is wider when the outcome uncertainty is lower.

Next, we quantify again *CCE* as a possible measure of SoA. *CCE* diminishes with outcome uncertainty even when compared at the same level of causal prior (Fig. 6a). Hence, *CCE* explicitly depends on the outcome uncertainty. When plotted as functions of temporal disparity, with the specific causal priors obtained for each outcome uncertainty condition, the peak values of *CCE* noticeably differ across the uncertainty conditions (Fig. 6b). This is because of the different values of the outcome uncertainty σ_*O*_ but also partly because of the different values of the causal prior. In all conditions, *CCE* falls off with the disparity of sensory inputs from the prior mean, |*τ*_*O*_ − *τ*_*A*_ − *μ*_*AO*_|. This fall off is milder when the uncertainty is lower. These results clearly manifest again three basic requirements of *CCE*: (A) the consistency of sensory inputs with the causal prior; (B) strong prior belief in causality; and (C) reliable sensory inputs.

**Figure 6.**
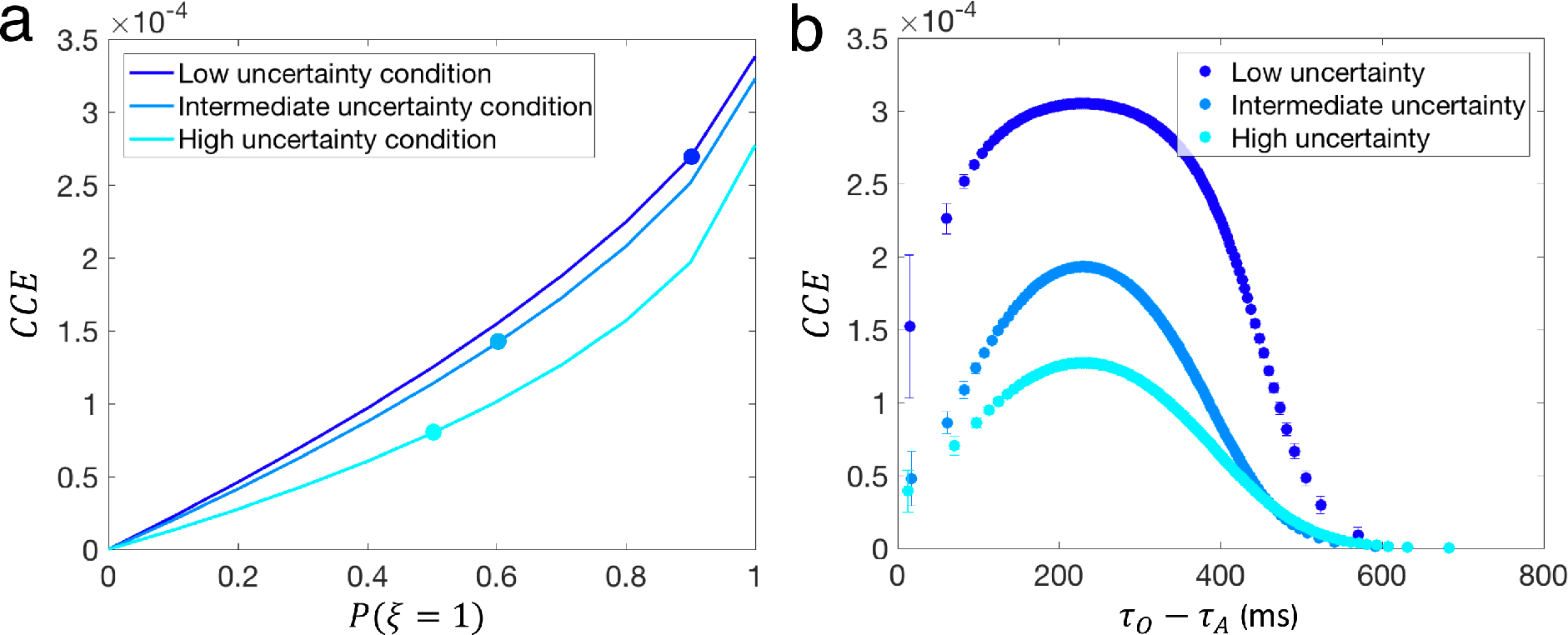
The Bayesian model prediction of *CCE* as a function of **(a)** the causal prior and **(b)** the temporal disparity *τ*_*O*_ − *τ*_*A*_. **a**| The different effects of the causal prior on *CCE* across the three conditions is evident even with equal causal priors, which means that *CCE* depends on outcome uncertainty. **b**| When plotted as functions of the temporal disparity *τ*_*O*_ − *τ*_*A*_, given the condition-dependent causal priors (marked by the colored dots in **(a)**), *CCE* falls off with the disparity of sensory inputs from the prior mean, |*τ*_*O*_ − *τ*_*A*_ − *μ*_*AO*_|, faster when the outcome uncertainty is higher.

## Discussion

We formalize SoA by drawing parallels from a Bayesian inference of the *ventriloquism effect* that estimates a common cause behind its multisensory integration. Our Bayesian model integrates the action-outcome signals, compares them with the prior expectation, and infers the causality between them as well as the timing of these sensory signals. Our model could concisely reproduce the intentional binding experiments by Haggard et al.’s and Wolpe et al. The temporal binding or repulsion phenomena are explained by the compromise between the noisy sensory observations and the prior belief of the action-outcome timing. Importantly, our Bayesian model predicts that the perceptual binding is generally trial-dependent, and it must be correlated with the estimated causality 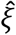 between the action and outcome. This prediction can be tested, when the probability *P_c_* for detecting causality is not close to 0 or 1, by examining if the distribution of action-outcome intervals is bimodal and if the intervals correlate with the reported causality between the action and outcome. In this work, we focus on the timing to investigate the intentional binding effects but the mathematical elucidations of our model can permit other modalities (e.g., visual or haptic) and structural properties (e.g., inter alia, location, size, shape and texture).

In addition, we theorize SoA as *the confidence in causal estimate*: *CCE*. *CCE* is high when the action-outcome timing is consistent with the causal prior, the causal prior is strong, and the action and outcome signals are reliable. This notion is consistent to what have been propounded as demonstrations of SoA: SoA arises from the causal relation between performed actions and their consequences^4,19,22,23^ and from the integration of different agency cues whose individual influences are determined by their reliability^14,15,24-27^. Hence, we posit *CCE* to be a plausible measure of SoA.

Specifically, we postulate *CCE* fits the notion of a pre-reflective, implicit *feeling of agency* (FoA). Synofzik and colleagues^4,25^ provide a compelling account of such feeling: FoA is best accounted for by multimodal weighting and integration of different agency cues, and consists of an automatic registration of whether an action or sensory event is caused by the self or not. They posit FoA is nothing other than first-person in that the self is implied, hence, no external attribution (e.g., to TMS that caused the action) is possible. In the event that there is a feeling of exogenous causation, this will be overwritten by an explicit, interpretative judgment of agency (JoA) based on contextual beliefs or rationalizations. Similarly, the analytical expression of *CCE* shows that it is a multimodal weighting and integration process that lies at the center of obtaining a Bayesian causality inference. Furthermore, *CCE* itself does not attribute causality to any external agent, such as in the case of strong causal prior for TMS-induced movements. The judgement of the causality, 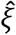, is then made based on the posterior ratio *r* that compares *CCE* with the confidence in the acausal estimate. Perceptual timing in our model simply reflects the sensory signals if the causality is not detected 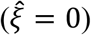, whereas they are overwritten by the influence of the prior if the causality is detected 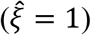. For example, in the involuntary condition of Haggard et al., the estimated action and outcome timing by the model repulse reflecting the judgment of the causality. A compelling speculation in Haggard et al.’s paper^6^ suggests this notion: the repulsion in the involuntary condition “reflects a mental operation to segregate, and thus to discriminate, pairs of events that cannot plausibly be linked by our own causal agency” (p. 384). We suggest such mental operation fits the notion of JoA, as quantified by the time shifts in equation (5) with the detected causality, and the peculiar feeling of causation by the involuntary movement to be FoA, quantified by *CCE*.

Following the above explanation, our theory therefore has a different take of Haggard et al.’s binding effect that requires intentionality. *CCE* argues that the judgment of the causality is central to the perceived temporal binding, consistent with current evidence that competes with the intentional account: the temporal binding is actually *causal*, not intentional^22^. For example, our model judges the causation of the tone even by the TMS-induced action in the involuntary condition. Hence, our Bayesian model predicts this unintended causality. Furthermore, our Bayesian model predicts that the action-outcome timing shifts toward the prior belief, 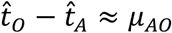, when the causality is perceived irrespective of the nature of the action, whether self-generated (i.e., the voluntary condition) or unintended (i.e., the involuntary condition). We suggest that the repulsion happened in the involuntary condition because of the proprioceptive noise (large σ_*A*_) that characterizes the TMS. The large proprioceptive noise may be caused by the internal prediction error resulting from unintended proprioceptive signal^28,12^ or the large clicking sound of TMS. In this sense, intentionality is just one factor that influences *CCE* and perceptional shift in our model. We predict that an experimental manipulations that reduces σ_*A*_ would increase perceived SoA even for unintended artificial actions. The prediction is therefore distinct from what was previously considered and can therefore serve as testable prediction for future experiments on causal agency.

Our theory also has a different take of Wolpe et al.’s binding effect. Wolpe and colleagues showed intentional binding as cue integration with uncertainty in outcome signals. They speculated that action and outcome bindings are driven by two distinct mechanisms: action binding is predicted by cue integration, but outcome binding supports the predictive pre-activation hypothesis^29^, i.e., the neural representation of the sensory outcome is activated prior to it. Hence, the outcome signals are perceived faster with less jitter than when it is not predicted to occur after the action. This could explain why the subjects’ timing estimations are largely erroneous in the baseline condition and why the outcome binding is greater than the action binding. Our theory, although qualitative, explains both action and outcome bindings by a single Bayesian cue integration mechanism. Our model explains that the magnitudes of the action and outcome perceptual shifts, (*τ*_*O*_ − *τ*_*A*_ − *μ*_*AO*_, are influenced primarily by the ambiguity of the outcome sensory signals, 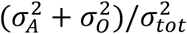, and also in part by the strength of the causal prior that diminishes with outcome uncertainty.

The intentional binding paradigm has also been used to study pathological sense of agency^30-32^. Patients with schizophrenia tend to have much stronger temporal binding than healthy volunteers. Moreover, unlike healthy volunteers, their temporal binding of action timing does not depend on the probability of the outcome tone presentation^32^. These results are explained by our Bayesian model by assuming that schizophrenia patients cannot easily adapt their abnormally strong belief in causality (i.e., too large *P*(*ξ* = 1)) and the uncertainty in the outcome (i.e., σ_*O*_). Another important point is that, unlike healthy volunteers, patients with schizophrenia exhibit temporal binding of action timing that depends on the presence or absence of the outcome. It will be an interesting future study to model this result by explicitly incorporating the probabilistic occurrence of the outcome in our Bayesian model.

In summary, we posit that since the Bayesian cue integration is primarily precision-dependent so is our theory of SoA. Our model predicts that if the uncertainty of the sensory input signals could be maintained small, even unintended causal action may give rise to high *CCE* (hence, strong SoA) - hence, our notion of precision-dependent casual agency. We posited the precise estimation that gives rise to SoA encapsulates consistency in the perceived action-outcome effect, the prior belief of the causation of the outcome by the action, and the reliability of the perceived sensory signals. This theory may shed light on the mechanism of reduced SoA in psychosis, the understanding of the difference between FoA and JoA, and the design of prosthetic devises that heighten SoA.

## Acknowledgement

This study was supported by Brain/MINDS from AMED under Grant Number JP18dm020700 and JSPS KAKENHI Grant Number JP18H05432.

## Methods

### Analytical Expressions for the Bayesian Estimates

The MAP estimate (equation (3)) of the Bayesian observer has a simple analytical expression. The MAP estimation is computed based on the posterior probability *P*(*t_A_*, *t_O_*, *ξ*|*τ*_*A*_, *τ*_*O*_) = *P*(*τ*_*A*_, *τ*_*O*_, *t_A_*, *t_O_*, *ξ*)/*P*(*τ*_*A*_, *τ*_*O*_), where the peak location only depends on the joint distribution *P*(*τ*_*A*_, *τ*_*O*_, *t_A_*, *t_O_*, *ξ*) in the numerator. The joint distribution is decomposed as *P*(*τ*_*A*_, *τ*_*O*_, *t_A_*, *t_O_*, *ξ*) = *P*(*τ*_*A*_|*t*_*A*_)*P*(*τ*_*O*_|*t*_*O*_)*P*(*t*_*A*_, *t*_*O*_|*ξ*)*P*(*ξ*), where the conditional distributions for action and outcome are 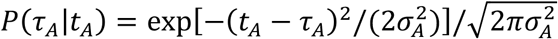 and 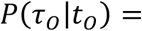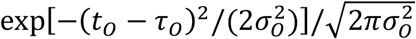, respectively, and the prior distribution is

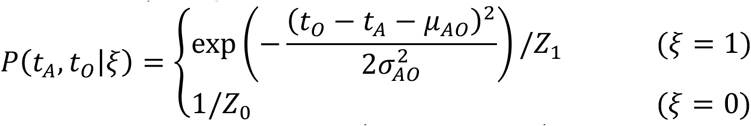

with normalization constants 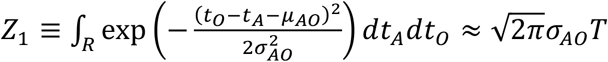 and *Z*_0_ ≡ ∫_*R*_*dt*_*A*_*dt*_*O*_ = *T*^2^.

We separately compute the peak location 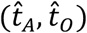 for the causal case *ξ* = 0 and the acausal case *ξ* = 0 and, then, compare these two peaks. In the acausal case, because *P*(*τ*_*A*_|*t*_*A*_) and *P*(*τ*_*O*_|*t*_*O*_) take the maximum values at *t*_*A*_ = *τ*_*A*_ and *t*_*O*_ = *τ*_*O*_, respectively, the location of the acausal peak is 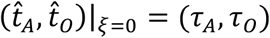 and the peak value is 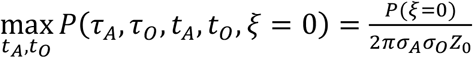. In the causal case, the peak of the joint distribution is found by minimizing a quadratic function. The peak location is given by 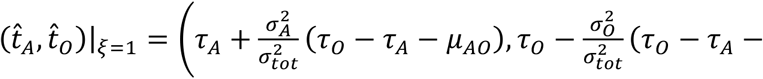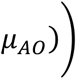, where 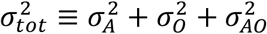 is the total variance, and the peak value is computed as 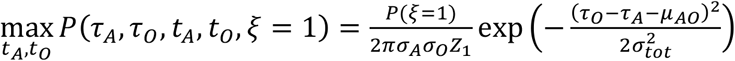. We define the log-ratio of the posterior peaks for *ξ* = 1 and *ξ* = 0 by

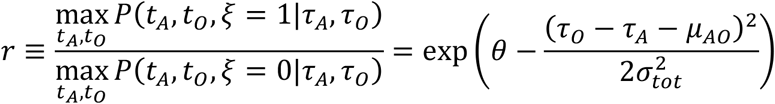

with 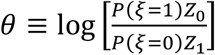. If *r* > 1, the MAP estimate is given by 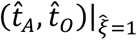 and 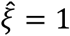, which predicts perceptual shifts. If *r* < 1, the MAP estimate is given by 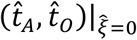 and 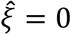, which predicts no perceptual shifts. The probability for detecting causality (i.e., 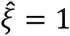) is also easily computable because *τ*_*O*_ − *τ*_*A*_ − *μ*_*AO*_ is distributed according to the Gaussian distribution 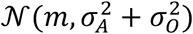 with 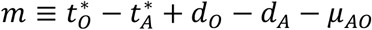. Hence, the causality is detected if 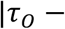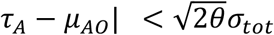, and this happens with probability

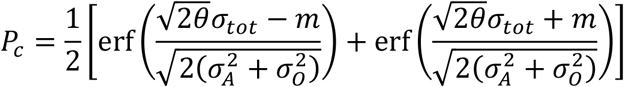

Next, we evaluate the confidence in the causal MAP estimation 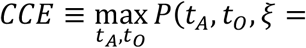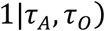, which comprises the numerator of the ratio *r*. To quantify this confidence, we need to first evaluate *P*(*τ*_*A*_, *τ*_*O*_) = *P*(*τ*_*A*_, *τ*_*O*_, *ξ* = 1) + *P*(*τ*_*A*_, *τ*_*O*_, *ξ* = 0) with

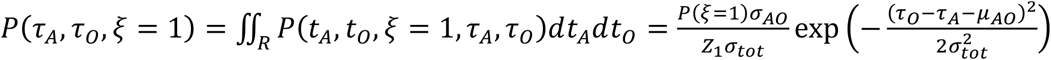

and

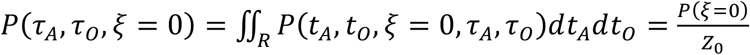

Combining these expressions together, we obtain

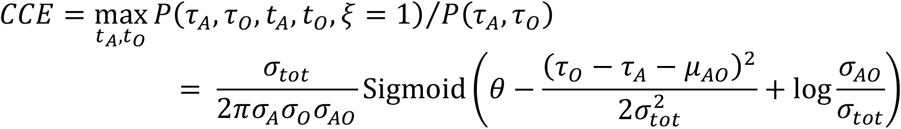

where Sigmoid(*x*) = 1/(1 + *e*^−*x*^) is the sigmoid function.

### Validation of Bayesian Estimates

The principal measure of intentional binding is the mean perceptual shift of temporal awareness of action and sensory outcome. A *perceptual shift* is the change in the subjective estimation of action or outcome timing from the baseline (*Bsln*) to the operant (*Oprnt*) condition, which is computed as

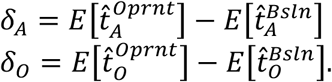

A *positive* shift informs the perception of timing shifted later in time, and a *negative* shift informs the perception of timing shifted earlier in time.

With the mean action and outcome perceptual shifts, δ_*A*_ and δ_*O*_, we can obtain the *model estimation error*, i.e., the difference of our model’s (*Model*) prediction of the action or outcome perceptual shift to the corresponding perceptual shift reported in the experiments (*Exp*). We obtain this as

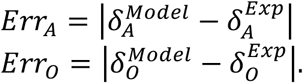

### Simulation Details

Table 1 lists all the parameters of our Bayesian model. We performed different simulations to reproduce the action and outcome perceptual shifts that were reported in the experiments, and explain their underlying psychophysical mechanisms in Bayesian terms. We generated 35,000 instances of *τ*_*A*_ and *τ*_*O*_ pairs for each experimental condition. To determine the optimal model parameter values, we performed *model fitting*: (*i*) a set of possible model parameter values are used to simulate and obtain a prediction dataset, (*ii*) the model estimation error (explained above) is computed to determine the difference of our model’s predictions to the reported empirical results, and (*iii*) the model parameters that best minimized this error are selected.

#### Simulations 1|

The objective was to determine the values of the operant parameters *μ*_*AO*_ and σ_*AO*_ that best estimate the results reported by Haggard, Clark & Kalogeras^6^. We tested using the baseline parameters in Table 1-Set A and different combinations of *μ*_*AO*_ and σ_*AO*_. We assumed initially that the action always causes the outcome, i.e., *P*(*ξ* = 1) = 1. We later dropped this assumption to test the effect of different causal priors.

For each *μ*_*AO*_ and σ_*AO*_ pair, we obtained the model estimation errors for the reported action and outcome perceptual shifts listed in Table 2-Set A. We took the average of the model estimation errors for the voluntary, involuntary and sham conditions to obtain a single model estimation error. We looked at the model estimation errors for the (a) action perceptual shifts only, (b) outcome perceptual shifts only, and (c) action-outcome perceptual shifts. After testing different *μ*_*AO*_ and σ_*AO*_ pairs, our results showed the best estimates of the model to be at *μ*_*AO*_ = 230 and σ_*AO*_ = 10. Furthermore, we observed the perceptual shift in action timing alone was sufficient to indicate the best estimates of the model.

**Table 2.** Reported perceptual shifts in action and outcome temporal awareness

| **Judged event (operant condition)** | *Mean estimation error (ms)* | *Mean perceptual shift (ms)* |
| --- | --- | --- |
| • *Set A*: Reported by Haggard, Clark & Kalogeras<sup>6</sup> |  |  |
| Voluntary action, then | 21 | 15 |
| tone | -31 | -46 |
| Involuntary action, then | 56 | -27 |
| tone | 46 | 31 |
| Sham TMS, then | 25 | -7 |
| tone | 7 | -8 |
| • *Set B*: Reported by Wolpe, Haggard, Siebner & Rowe<sup>20</sup> |  |  |
| Action, then | 31 | 39 |
| low uncertainty tone | -16 | -51 |
| Action, then | 23 | 31 |
| intermediate uncertainty tone | -19 | -65 |
| Action, then | 24 | 32 |
| high uncertainty tone | -10 | -105 |

#### Simulations 2|

The objective was to obtain the causal prior probability that yield the best estimates of Haggard et al.’s results. With *μ*_*AO*_ = 230 and σ_*AO*_ = 10, we tested for *P*(*ξ* = 1) in the range 0 to 1 with increments of 0.1. We used the same pairs of *τ*_*A*_ and *τ*_*O*_ from simulations (1), together with the Table 1-Set A baseline parameters, when we carried out our simulations. We computed once again the model estimation errors for the empirical results listed in Table 2-Set A. We selected the *P*(*ξ* = 1) that best minimized the estimation errors for the voluntary, involuntary and sham conditions. Table 1-Set C includes the parameters that yielded the best model estimates. Fig. 1 shows the action and outcome perceptual shifts, as well as the intervals between perceptual shifts, which were obtained by our Bayesian model using these parameters.

#### Simulations 3|

The objective was to reproduce the perceptual shifts reported by Wolpe, Haggard, Siebner & Rowe^20^, listed in Table 2-Set B. We generated another set of 35,000 *τ*_*A*_ and *τ*_*O*_ pairs using, this time, the baseline parameters listed in Table 1-set B. We performed simulations and tested using *μ*_*AO*_ = 230, σ_*AO*_ = 10, and *P*(*ξ* = 1) in the range 0 to 1 with increments of 0.1. We did not perform additional simulations to redetermine *μ*_*AO*_ and σ_*AO*_ since our aim is to reproduce qualitatively all the experiments with the same *μ*_*AO*_ and σ_*AO*_ as possible in order to have simple yet consistent explanations by our Bayesian model. Although we did not modify here *μ*_*AO*_ and σ_*AO*_, our analyses and results can show that their effects can be predicted and explained by our model. The model estimation errors once again indicate the estimates of action perceptual shifts led to the best estimates of the model. We list under Table 1-Set D the *P*(*ξ* = 1) that yielded the best estimates of the model for the low, intermediate and high uncertainty tone conditions. We show in Fig. 5a the action and outcome perceptual shifts, and intervals between shifts, predicted by our Bayesian model for this experimental setup.

#### Simulations 4|

The objective was to determine the influence of the causal prior and the temporal difference *τ*_*O*_ − *τ*_*A*_ (that varies in every trial) on the various predictions of our Bayesian model for Haggard et al.’s experimental setup. We used the model parameters and *τ*_*A*_ and *τ*_*O*_ pairs from simulations (1) and (2). We obtained our Bayesian model’s predictions of the intervals between action and outcome perceptual shifts, binding and repulsion effects, action-outcome timing interval, 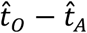, in the baseline and operant conditions, and *CCE*. The results are shown in Figs. 2, 3 and 4.

#### Simulations 5|

The objective and target results were the same as simulations (4), but using this time the model parameters and *τ*_*A*_ and *τ*_*O*_ pairs from simulations (3) to account for Wolpe et al.’s experimental setup. The resulting plots are shown in Figs. 5 and 6.

